# Comparative Transcriptome Analysis Unveils Mechanisms of Salt Tolerance in Bluebunch Wheatgrass

**DOI:** 10.64898/2026.08.04.742830

**Authors:** Yuanyuan Ji, Zhengping Wang, Raju Chaudhary, Sampath Perumal, Pierre Hucl, Bill Biligetu, Andrew G. Sharpe, Lingling Jin

## Abstract

Bluebunch wheatgrass (*Pseudoroegneria spicata*) exhibits substantial variation in its response to salt stress, making it a valuable model for studying salinity-tolerance mechanisms for use in crop improvement. In this study, we identified two *P. spicata* genotypes with contrasting responses to salt stress: the tolerant W6 56551, which maintained growth with green foliage under saline conditions, and the susceptible PI693916, which exhibited severe leaf chlorosis and stunted growth. To better understand the molecular basis of salt tolerance in bluebunch wheatgrass, we conducted RNA-sequencing at 0, 1, and 4 days (D0, D1, and D4) after salt treatment at 160 mM level to examine changes in gene expression of salt-tolerant and salt-susceptible genotypes. Comparative analysis across time points identified 6,154 and 1,086 differentially expressed genes (DEGs) at D4 and D1 in PI693916, and 4,638 and 3,302 DEGs at D4 and D1 in W6 56551, respectively, relative to control (D0). Functional analysis of these DEGs showed that the salt-tolerant genotype displayed an early and broad transcriptional reprogramming, including induction of photosynthesis, carbon metabolism, and flavonoid biosynthesis pathways, whereas the salt-susceptible genotype exhibited delayed and less coordinated responses, with enrichment of cyanoamino acid metabolism and repression of antioxidant-associated pathways. Notably, calcium signaling, ion transporter regulation, and osmolyte biosynthesis genes showed contrasting expression between genotypes, highlighting distinct strategies for ionic and osmotic homeostasis. Collectively, these results demonstrate that salt tolerance in *P. spicata* is associated with rapid metabolic adjustment, enhanced photosynthetic stability, and differential regulation of ion transport and osmoprotectant pathways.

## 1 Introduction

More than 20% of cultivated land and 50% of irrigated land are affected by salinization or at risk of salinization globally^1,2^, causing plant growth inhibition, disrupted development, metabolic adaptations and ion sequestration or exclusion^3,4^. Bluebunch wheatgrass (*Pseudoroegneria spicata*) is a highly palatable perennial grass widely used for livestock and wildlife. Its observed tolerance to salinity, broad ecological adaptability, and substantial genetic diversity^5–7^ indicate that natural variation in salinity response may exist within the species. This variation provides a valuable resource for investigating the molecular mechanisms underlying its potential capacity for salinity tolerance. As soil salinity becomes one of the most detrimental environmental factors limiting plant growth and development, enhancing salinity tolerance is highly desirable for improving forage production. Therefore, a deeper understanding of the mechanisms underlying salinity tolerance in bluebunch wheatgrass will be significant to the generation of salt-tolerant cultivars.

Salt stress is a multifaceted condition impacting plant growth and physiology. It induces three major types of stress: osmotic, ionic, and secondary stress ^8^. Excess soluble salts lower the water potential at the root surface, reducing water availability and limiting the uptake of water by plants, therefore triggering *osmotic stress* ^9^. *Ionic stress* results from the excess salt ions accumulated in leaves, disrupting metabolic pathways through disabling catalytic enzyme activities ^10^. *Secondary stress*, amplified by both osmotic and ionic stress, triggered the overproduction of reactive oxygen species (ROS) such as hydroxyl radical, hydrogen peroxide, and superoxide anions, which in turn damaged cellular structures and essential macromolecules^11,12^.

At the molecular level, salt stress triggers several well-characterized signaling pathways. For example, one of the earliest responses is a rapid rise in cytosolic Ca^2+^, which acts as a primary signal for stress perception^13^. The calcineurin B-like proteins and CBL-interacting protein kinases (CBL–CIPK) network is also triggered by salt stress. CBL-CIPK is among the best-studied calcium-decoding systems in plants ^14^ and has been characterized in multiple crops, including rice^15^ and wheat^16^. In this pathway, CBLs bind Ca^2+^ and interact with CIPKs, and their phosphorylation states translate calcium signatures into downstream regulatory outputs that shape transcriptional responses ^17–19^. Beyond calcium signaling, ion homeostasis, particularly the movement of Na^+^ and Cl^-^, is central to salt tolerance. Na^+^ uptake is mediated largely by nonselective cation channels (NSCCs), low-affinity cation transporters (LCT1), cation–chloride cotransporters (CCCs), and high-affinity K^+^ transporters (HKTs). Cl^-^ homeostasis involves transporters such as chloride channels (CLC), all of which help maintain ionic balance under saline conditions^20,21^.

These signaling and transport pathways have been extensively studied in Arabidopsis^22^ and several grass crops, including barley^23,24^ and wheat^25,26^. However, their roles in bluebunch wheatgrass remain largely unexplored. Whether these conserved pathways similarly contribute to salinity acclimation in *P. spicata* is still unknown. Moreover, one of the key mechanisms for osmotic adjustment is the accumulation of compatible solutes, such as proline, glycine betaine, sorbitol, and mannitol. These compounds are thought to play a protective role under stress conditions, functioning as antioxidants to mitigate the harmful effects of salt stress^27^. It is important to understand how the metabolism of these solutes is regulated at the transcriptional level.

In this study, we identified two phenotypically distinct *P. spicata* genotypes in response to salt stress, revealing one genotype (W6 56551) as salt-tolerant and the other (PI693916) as salt-susceptible. Comparative transcriptome analysis using RNA-seq on both genotypes under salt stress across multiple time points revealed dynamic transcriptional responses. The observed variation in transcriptome profiles between these genotypes provides a strong foundation for investigating the molecular mechanisms and regulatory networks underlying salt tolerance. Specifically, our objectives are to (1) identify differentially expressed genes (DEGs) and perform the enrichment of KEGG pathways to map DEGs to molecular networks, (2) characterize DEGs associated with Ca^2+^ signaling and ion transporters and evaluate their potential roles in salt tolerance, and (3) examine distinct patterns of secondary metabolism regulation associated with contrasting tolerance.

## 2 Materials and Methods

### 2.1 Plant Materials and Treatments

*P. spicata* plants were grown in a growth chamber for three weeks (24 °C, 16/8 h light/dark). 21-day-old seedlings of W6 56551 and PI693916 were treated with 100 mL of 160 mM NaCl solution for seven days for salt stress. The leaf samples were collected at 0-day (D0), 1-day (D1), and 4-day (D4) with three independent biological replicates, respectively. All materials collected were flash-frozen in liquid nitrogen and stored in a -80 °C freezer until required. Seeds of both genotypes were obtained from Genebank at the National Plant Germplasm repository, United States Department of Agriculture in Ames, Iowa.

### 2.2 RNA-seq Sequencing

Total RNA was extracted from 50 mg of materials using Illumina stranded mRNA Prep Kit (Illumina). RNA-seq libraries were prepared using SMRTbell prep kit 3.0 (PacBio) and sequenced in Novaseq 6000 platform (2 × 150 bp). A total of 721,859,179 raw reads is generated. The raw paired-end reads from RNA-seq were subjected to quality trimming using Trimmomatic v.0.39 with default settings ^28^, generating 683,866,591 clean reads (TableS1).

### 2.3 Identification of Differentially Expressed Genes (DEGs)

Clean reads were aligned to *P. spicata* (PI635993) reference genome ^29^ using STAR v2.7.6 ^30^ with default parameters. Gene expressions were quantified using RSEM (v1.3.3) ^31^. The expression level was normalized in fragments per kilobase million (FPKM) values and counts per million (CPM) for expression analysis. DESeq2 (v1.42.0)^32^ was used to identify significant DEGs with false discovery rate (FDR) *<* 0.1 and log fold change (LFC) ≥ 2. Multidimensional scaling plot (Figure S1B), principal component analysis (Figure S2B), and the distribution of expression level in each replicate are performed using integrated Differential Expression and Pathway (iDEP) analysis web application^33^ with raw read counts as input.

### 2.4 KEGG Pathway Enrichment

Protein sequences of *P. spicata* reference genome were first annotated to Entrez gene ID using KOBAS (v3.0) ^34^ with *Brachypodium distachyon* as a reference. Entrez gene IDs corresponding to the DEGs were extracted from each comparison and used as input. The enrichment was conducted against the *Brachypodium distachyon* reference pathway database (KEGG organism code:bdi) using clusterProfiler (v4.16.0) ^35^. Enrichment significance was determined with a *p*-value cutoff of 0.5. The top 10 pathways were visualized using dot plots generated using ggplot2 in R ^36^.

## 3 Results

### 3.1 Contrasting Salt Tolerance Phenotypes in *P. spicata* Accessions

Previous evaluations identified several *P. spicata* accessions as exhibiting marginal salinity tolerance ^5^. Moreover, our recent study suggests that the species harbors substantial natural genetic variation^6^. Through an initial large-scale screening of 145 genotypes (TableS2), we revealed substantial variation in salt tolerance within the *P. spicata* populations. To further elucidate the physiological and molecular mechanisms underlying salt resillience in *P. spicata*, we selected two phenotypically divergent accessions for comparative evaluation: W6 56551 (Utah) and PI693916 (Washington). These accessions were subjected to a rigorous high-salinity regime of 16 dS/m, a threshold at which the majority of related graminaceous species exhibit acute growth inhibition^37^.

After one week of salt treatment (Methods), the two accessions displayed strikingly different phenotypes. W6 56551 maintained vigorous growth, characterized by elongated green leaves and continued tiller development (Figures 1A and 1B). In contrast, PI693916 showed clear symptoms of salt sensitivity, including leaf chlorosis, reduced leaf elongation, and minimal branching (Figures 1A and 1B). These contrasting responses indicate that W6 56551 possesses substantially greater salt tolerance relative to PI693916, highlighting the presence of exploitable natural variation for salinity response within *P. spicata*.

**Figure 1.**
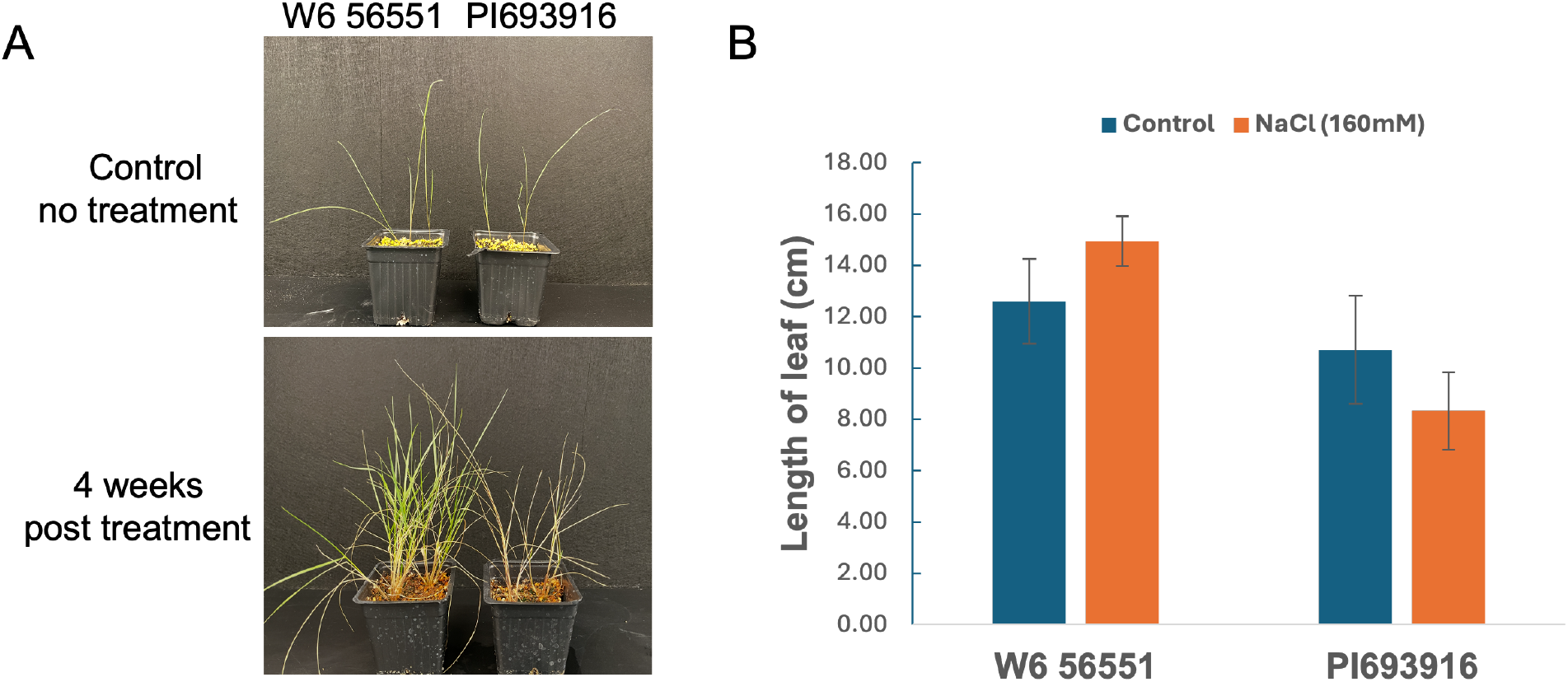
Phenotypic response of bluebunch wheatgrass (W6 56551 and PI693916) to salt stress treatment (160 mM NaCl). Plants were treated with 100 mL of 160 mM NaCl solution for one week. Control plants (top) were photographed at day 0, prior to treatment (no treatment). Salt-stressed plants were photographed 4 weeks after treatment.

### 3.2 Transcriptomic Profiling of Differentially Expressed Genes Under Salt Stress Treatment

RNA-sequencing was performed on eighteen leaf samples of *P. spicata* (2 genotypes × 3 time points × 3 replicates), yielding approximately 11.08 Gb of data per sample (199.52 Gb in total). The average mapping rate to the reference genome was 99.98% (TableS2), and 36,450 of 36,559 annotated genes showed detectable expression. These results indicate that the sequencing data are reliable and the reference genome is well suited for downstream analyses. The overall gene expression level across treatments and genotypes, each with three biological replicates, were comparable based on transformed CPM values (Figure S1A). The multidimensional scaling analysis (MDS) of read counts from all samples revealed that most biological replicates clustered tightly, with slight dispersion observed in PI693916 samples at Day 1 and 4 post-treatment (Figure S1B).

Differentially expressed genes were analyzed across seven comparisons: D4 vs. D1, D1 vs. D0, and D4 vs. D0 in both genotypes, as well as PI693916_D0 vs. W6 56551_D0. DEGs were identified using thresholds of FDR *<* 0.1 and LFC ≥ 2. In PI693916, DEG counts were highest in D4 vs. D0 (2,710 up-regulated, 3,444 down-regulated) and D4 vs. D1 (1,841 up-regulated, 3,390 down-regulated), but lower in D1 vs. D0 (762 up-regulated, 324 down-regulated) (Figure 2A). In contrast, W6 56551 showed fewer DEGs in D4 vs. D1 (697 up-regulated, 2,107 down-regulated) and D4 vs. D0 (1,997 upregulated, 2,641 down-regulated), but more in D1 vs. D0 (1,453 up-regulated, 661 down-regulated) (Figure 2A). Notably, the between-genotype comparison (PI693916_D0 vs. W6 56551_D0) identified 5,512 up-regulated and 4,617 down-regulated genes, substantially more than any within-genotype comparison, indicating extensive baseline transcriptional divergence under non-stress conditions (Figure 2A).

**Figure 2.**
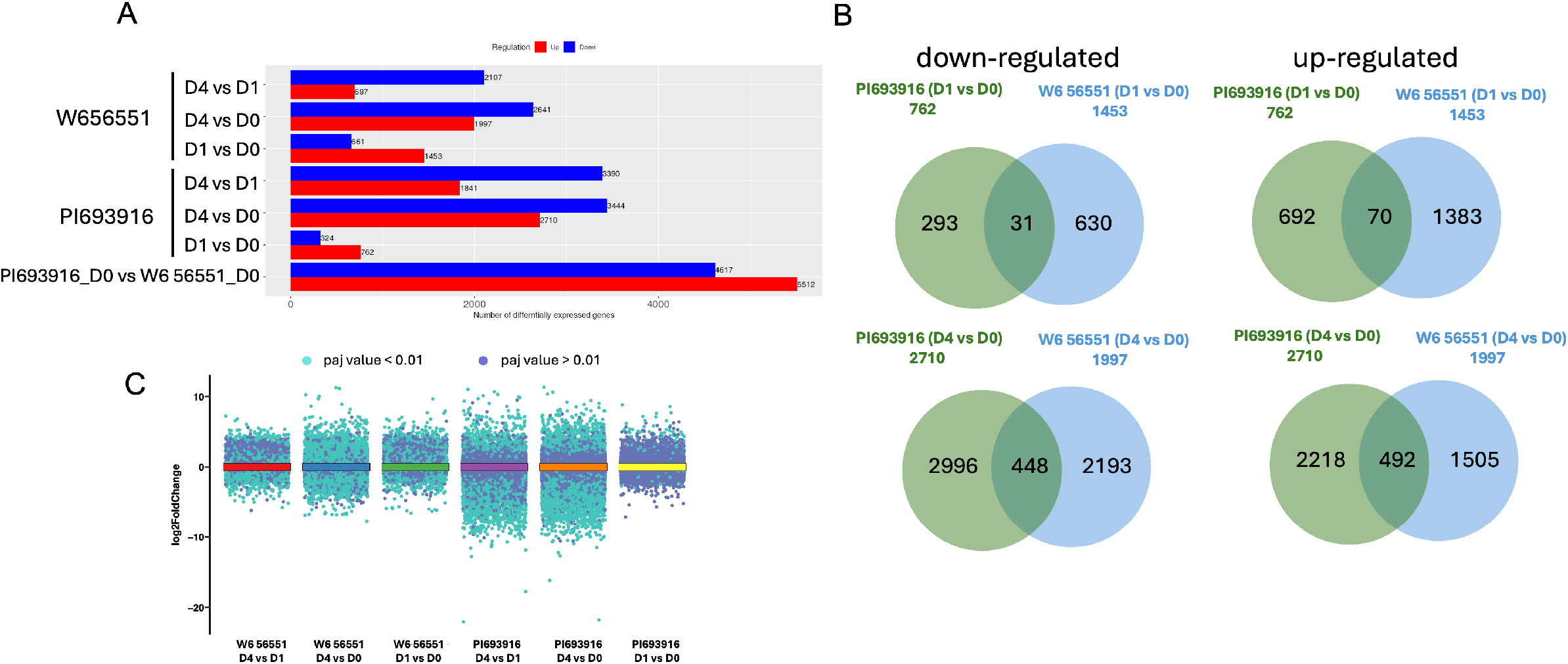
DEGs in genotype of W6 56551 and PI693916 under salt treatment at different time points. (A) The number of DEGs in different comparisons for each genotype. (B) Venn diagrams of DEGs between W6 56551 and PI693916 in 1 or 4 days after salt treatment. (C) Jitter plot showing the distribution of gene expression changes across six pairwise comparisons. The adjusted p value (*paj*) is calculated based on Benjamini-Hochberg procedure implemented in DESeq2. Genes were considered significantly differentially expressed with FDR *<* 0.1 and LFC *≥* 2.

Overlapping DEGs between D1 vs. D0 and D4 vs. D0 within each genotype are shown in Figure 2B, with expression patterns summarized in Figure 2C. Interesting, the proportion of overlapping genes remained consistent at approximately 3.2% for the D1 vs. D0 comparison in both genotypes (PI693916: 31/954, W6 56551: 70/2145). In contrast, this proportion increased significantly to over 7.9% in the D4 vs. D0 comparisons (PI693916: 448/5637, W6 56551: 492/4215). This results indicates an early pronounced transcription variation that converge as the duration of salt treatment progresses.

### 3.3 Early and Broader Transcriptional Reprogramming in Tolerant Genotype Reveals Mechanisms of Salt Response

To investigate the molecular mechanisms underlying salt stress tolerance, we performed functional enrichment analysis on DEGs from D1 vs. D0 and D4 vs. D1 comparisons in both genotypes. For early responses (D1 vs. D0), PI693916 showed minimal transcriptional response, with only two enriched pathways among the up-regulated DEGs (cyanoamino acid metabolism and ABC transporters) and no pathways were enriched among the down-regulated DEGs (Figure 3A). In contrast, W6 56551 exhibited a broader and robust early transcriptional re-programming on the first day of treatment. Pathways related to photosynthesis, carbon metabolism, oxidative phosphorylation, and flavonoid biosynthesis were significantly induced, as determined by adjusted *p*-values or gene ratios (Figure 3C). Meanwhile, pathways associated with amino acid metabolism, secondary metabolite biosynthesis, and plant–pathogen interactions were significantly down-regulated.

**Figure 3.**
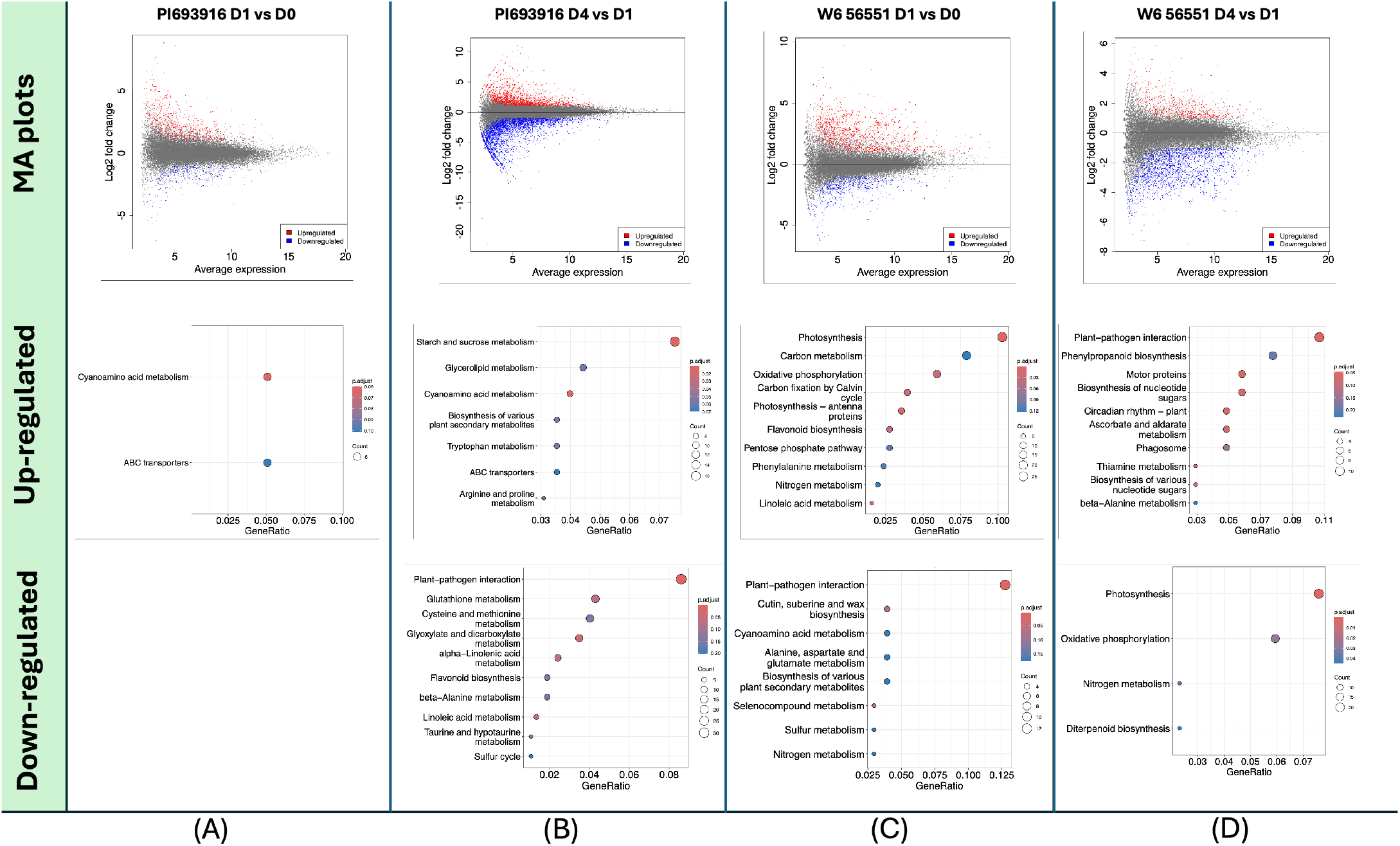
KEGG pathway enrichment of up/down-regulated DEGs in both genotypes. MA plots shows the overview of DEGs that up/down-regulated (top row).

In response to extended treatment (D4 vs. D1), PI693916 displayed delayed metabolic shifts, with significant up-regulation of starch and sucrose metabolism, cyanoamino acid metabolism, and glycerolipid metabolism (adjusted *p* value *<* 0.05 or gene ratio *>* 0.04), whereas pathways related to plant–pathogen interactions, glutathione metabolism, cysteine and methionine metabolism, glyoxylate and dicarboxylate metabolism, linolenic acid metabolism, and flavonoid biosynthesis were down-regulated (Figure 3B). Conversely, W6 56551 showed reverse regulation of pathways initially induced/suppressed at D1 including plant-pathogen interaction, photosynthesis, and oxidative phosphorylation, while maintaining upregulation of nucleotide sugar biosynthesis and ascorbate and aldarate metabolism in D4. Notably, flavonoid biosynthesis and linoleic acid metabolism, which were down-regulated in PI693916 at D4, were highly induced in W6 56551 at D1 (Figure 3B and 3D), suggesting their potential physiological roles in conferring early salt stress tolerance.

It is worth noting that nitrogen metabolism was enriched among both up-regulated and down-regulated genes in the D1 vs. D0 comparison of W6 56551. Despite sharing the same pathway category, the specific sets of genes involved differed between the two groups. The corresponding gene IDs and their annotated functions are provided in TableS6.

### 3.4 Divergence of Regulation on Calcium Signalling and Ion Transport Under Salt Stress Between Genotypes

The initial response to salt stress is a rapid induction of cytosolic Ca^2+^ levels, mediated by Ca^2+^ channels and interacting proteins such as calcineurin B-like (CBL) proteins ^38^. This Ca^2+^ signal subsequently coordinates multiple tolerance strategies such as antioxidant boosting, osmoprotectant accumulation, and ion uptake and transport^14,39,40^. To investigate the genetic basis of these processes in *P. spicata*, we identified the core components of the Ca^2+^ signaling network and key ion transporters. Our analysis revealed 8 calcineurin B-like protein genes (*CBL*), 25 non-specific CBL-interacting protein kinase genes (*CIPK*) (listed in TableS3). In addition, we also identified 99 ion/cation transporter genes including 11 glutamate-gated ion channels (also known as glutamate receptor-like, *GLR*), 3 cation transporter, 16 chloride channel, 15 potassium transporter, 22 sodium/hydrogen exchangers (*SOS*), 4 cyclic nucleotide-gated ion channel (*CNGC* s), and 28 non-selective cation channels (*NSCC* s) (listed in TableS4).

To better understand how these components contribute to differential salt tolerance, we examined the expression patterns in the salt-tolerant line (W6 56551) and the salt-sensitive line (PI693916). A clear divergence in transcriptional regulation was observed between the two lines. Specifically, in W6 56551, 2 of 8 *CBL* genes displayed high expression levels under control conditions, but downregulated with salt treatment, while 4 other *CBL* genes were induced by salt treatment either at D1 or D4. By contrast, their expression in PI693916 remained low and showed limited responsiveness to salt stress (Figure 4A). Interestingly, *CIPK* genes exhibited opposing regulatory trends between the two lines. In W6 56551, only 5 of 25 *CIPK* genes were induced by salt treatment at either D1 or D4, while the majority were strongly repressed or maintained at low expression levels (Figure 4A). In contrast, PI693916 showed induction of 14 *CIPK* genes following salt stress, indicating that *CIPKs* may play genotype-specific roles in salt stress signaling (Figure 4A).

**Figure 4.**
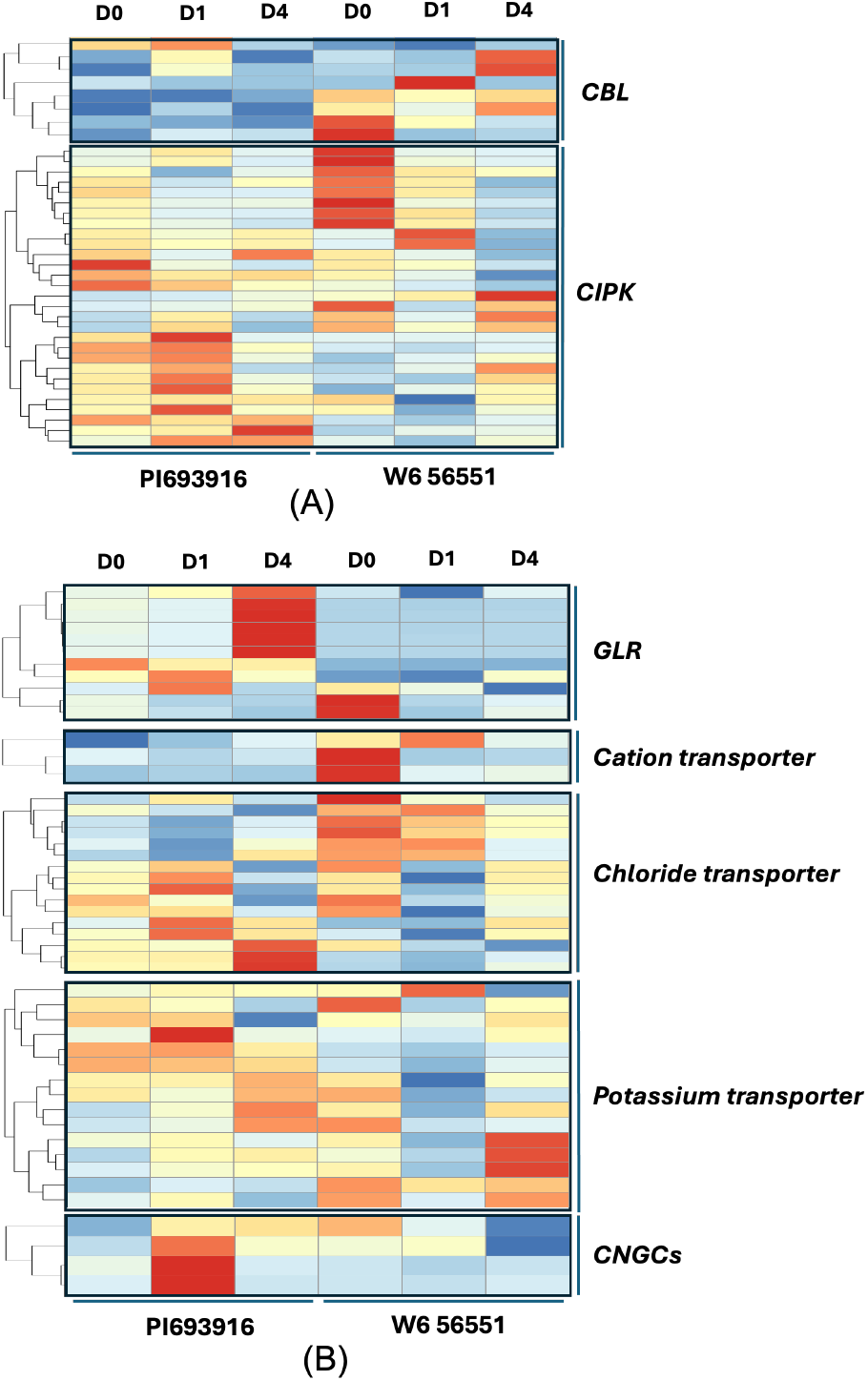
Transcript expression of genes associated with stress-responsive pathways. (A) Heatmap showing the expression of *CIPK* and *CBL* genes across all time points in W6 56551 and PI693916. (B) Heatmap showing the expression of *GLRs*, cation transporters, chloride channels, potassium transporters, and *CNGCs*.

We next examined the transcriptional expressions of ion/cation transporters in the two genotypes under salt stress. A similar expression pattern was observed for *GLR* and *CNGC* genes, which were strongly induced in the salt-sensitive line PI693916 but showed minimal change or repressed in tolerant line W6 56551 (Figure 4B). Cation transporter genes followed a distinct expression pattern, being downregulated in W6 56551 while maintaining low and unresponsive expression in PI693916 (Figure 4B). The downregulation of transporters in tolerant line suggests a reduced ionic fluxes or altered signaling activity in response to stress. Moreover, chloride transporters exhibited a more complex, and gene member-specific regulation pattern that differed fundamentally between genotypes. A subset of chloride transporter genes was specifically induced at D1 or D4 upon salt treatment in PI693916, while a partially overlapping subset of genes were strongly repressed in W6 56551 (Figure 4B), implying that non-redundant roles of individual chloride transporter are integrated into divergent ionic homeostasis strategies in two genotypes.

Most notably, potassium transporters in W6 56551 were rapidly and strongly repressed as early as D1 following salt treatment (Figure 4B), implying an early suppression of K^+^ uptake and transport processes. For *SOS* and *NSCCs* gene families, their expression displays a variable pattern, with several genes showing up-regulated under stress (D1 or D4), while a subset of members exhibits the opposite behavior (Figure 4B), suggesting distinct roles of each member within *SOS* and *NSCCs* families in response to salt conditions (Figure S3).

### 3.5 Potential Involvement of Proline and Glycine Betaine in Salt Resistance

Proline and Glycine betaine are two protective osmolytes under saline or stress conditions^27^. In plants, Proline is synthesized primarily from either Glutamate or ornithine, catalyzed by 1-pyrroline-5-carboxylate (P5C) synthetase (P5CS), reductase (P5CR), and ornithine aminotransferase (OAT), respectively. Meanwhile, Proline catabolism occurs with oxidation back to P5C and Glutamate via proline dehydrogenase (ProDH) and *δ*1-pyrroline-5-carboxylate dehydrogenase (P5CDH) ^41^ (Figure S3A). Glycine betaine synthesized from choline in a two-step reactions via a choline mono-oxygenase (CMO) and betaine aldehyde dehydrogenase (BADH) ^42^(Figure S3A). To explore their role in salt resistance in the W6 56551 line, we identified all homologous genes in *P. spicata*, with only one copy of each homolog detected; the gene IDs are listed in TableS5.

Regarding the Proline synthesis pathway genes, our results reveal that *GDH* and *PRODH* are the only two genes markedly induced by salt stress in W6 56551. In contrast, genes involved in Proline biosynthesis—*P5CS1, P5CS2, P5CR*, and *OAT* —were consistently down-regulated, suggesting a deceleration of Proline accumulation in this salt-tolerant genotype. Meanwhile, a notable pattern emerges for glycine betaine biosynthesis: *CMO* is transiently induced at D1 in W6 56551, whereas *BADH* expression is inhibited right after stress treatment. More interestingly, *CMO* and *BADH* genes show opposite regulation by salt stress in W6 56551, their expression levels in PI693916 are substantially lower than in W6 56551. This suggests that glycine betaine biosynthesis may play only a limited role in the salt stress response of the sensitive genotype.

## 4 Discussion

### 4.1 Transcriptional Landscapes and Metabolic Shift under Salt Stress

Salinity stress in plants is a dual-threat condition involving osmotic and ionic stress which limits water uptake and inhibits vital enzymatic processes simutaneously ^43^. In our analysis of *P. spicata* accessions PI693916 and W6 56551, we observed divergent phenotypic responses to salinity derived from fundamental difference in transcriptional timing and metabolic prioritization. A core set of genes involved in carbon metabolism including starch and sucrose metabolism and pyruvate metabolism were up-regulated in both lines (Figure 3), suggesting that central carbon metabolism is maintained to provide the energy necessary for adaptation. This increased carbon flow likely originates from enhanced glycolysis ^44^, aligning with previous observations in monocot crops where glycolysis and sucrose metabolism are co-induced under saline conditions^45–47^. However, a defining feature of the salt-tolerant W6 56551 is its rapid transcriptional response, showing a substantially higher number of regulated genes on the first day of treatment compared to PI693916, similar to patterns seen in contrasting cotton cultivars^48^. By promptly inducing photosynthesis and carbon metabolism at D1, W6 56551 minimizes the longterm damage of salt stress ^49^. This early metabolic activation is fundamental for tolerance, as seen in Arabidopsis^50^ and sweet sorghum^51^. Conversely, the sensitive PI693916 shows an upregulation of cyanoamino acid metabolism, which participates in the synthesis of cyanogenic glycosides and the production of hydrogen cyanide (HCN) ^52^. While HCN can serve as a nitrogen source^53,54^, its excessive production is toxic and impairs ATP production ^55^, potentially leading to the repression of flavonoid and linoleic acid metabolism—pathways usually associated with antioxidant defense and membrane stability^56,57^.

### 4.2 Calcium Signaling, Ion Homeostasis, and Osmolyte Strategy

Salt tolerance is further mediated by the calcineurin B-like protein (CBL) family and the highly conserved SOS signaling pathway (SOS1, CIPK24, and CBL4), which regulates Na^+^ efflux^58–62^. In W6 56551, the downregulation of *CBL7* homologs supports the notion that specific *CBL* members have antagonistic roles; while *CBL4/10* promote Na^+^ extrusion^63^, *CBL2/7* can negatively regulate tolerance by repressing H^+^-ATPase activity ^64^. The complexity of these interactions, such as the co-induction of *CBL4* and *CIPK6* in cucumber^65^ or the coordination of ion homeostasis by *CBL2/3* via several *CIPKs* ^66^, suggests that the CBL-CIPK network is highly context-dependent. Regarding transport, W6 56551 and PI693916 displayed opposite expression trends; most transporter genes were downregulated in the tolerant line but upregulated in the sensitive one (Figure 4). This is significant because transporters can function as either positive or negative regulators. For instance, loss-of-function mutants of *AtCNGC10* and *AtCNGC3* exhibit enhanced salt tolerance ^67–69^, whereas *athkt1;1* mutants accumulate less Na^+^ in roots ^70^, and *atglr3*.*4* mutants show increased sensitivity^71^. Efficient ion partitioning supports water balance and provides a favorable environment for osmolytes ^72^.

Efficient ion partitioning across tissues and cellular compartments not only supports water balance but also provides a favorable environment for organic solutes, which function as major osmolytes in the cytosol and organelles^72^. Genotypic differences in proline accumulation under salt stress have been extensively reported in crop species. In many cases, higher levels of free proline have been positively correlated with salt tolerance, and proline content has even been proposed as an index for assessing the tolerance potential of cultivars^73,74^. However, other studies have shown that salt-sensitive cultivars may accumulate significantly more proline than tolerant ones ^75–77^, suggesting that proline accumulation may sometimes reflect stress injury rather than adaptive tolerance. Based on our transcriptome analysis of genes involved in proline and glycine betaine metabolism, the expression patterns at both D1 and D4 indicate that, in W6 56551, salt tolerance is more strongly associated with glycine betaine metabolism than with proline accumulation.

## Acknowledgments

We extend our gratitude to Omics and Precision Analytics Laboratory (OPAL) and Data Management and Analytics (DMA) platform at GIFS for DNA sequencing and analysis. This work is funded by Natural Sciences and Engineering Research Council of Canada, the Agriculture Development Fund (ADF) under the Ministry of Agriculture of Saskatchewan and the Saskatchewan Cattleman’s Association.

## 5 Supplementary Results Availability

The supplementary tables in this study are available at Github (https://github.com/ifoo1213/RNA_seq-analysis-of-Bluebunch-Wheatgrass-under-salt-stress.git)

## 6 Author Contributions

Y. J., L.J. and A.S. conceptualized the study. Y.J. performed the salt stress experiment and RNA-seq analysis. Z.W. prepared the RNA-seq libraries. R.C. performed raw data preprocess. P.H and B.B. provide material. Y.J. wrote the first draft. B.B., L.J., and A.S. reviewed and revise the final version of the manuscript.

## 7 Supplementary Material

**Figure S1.**
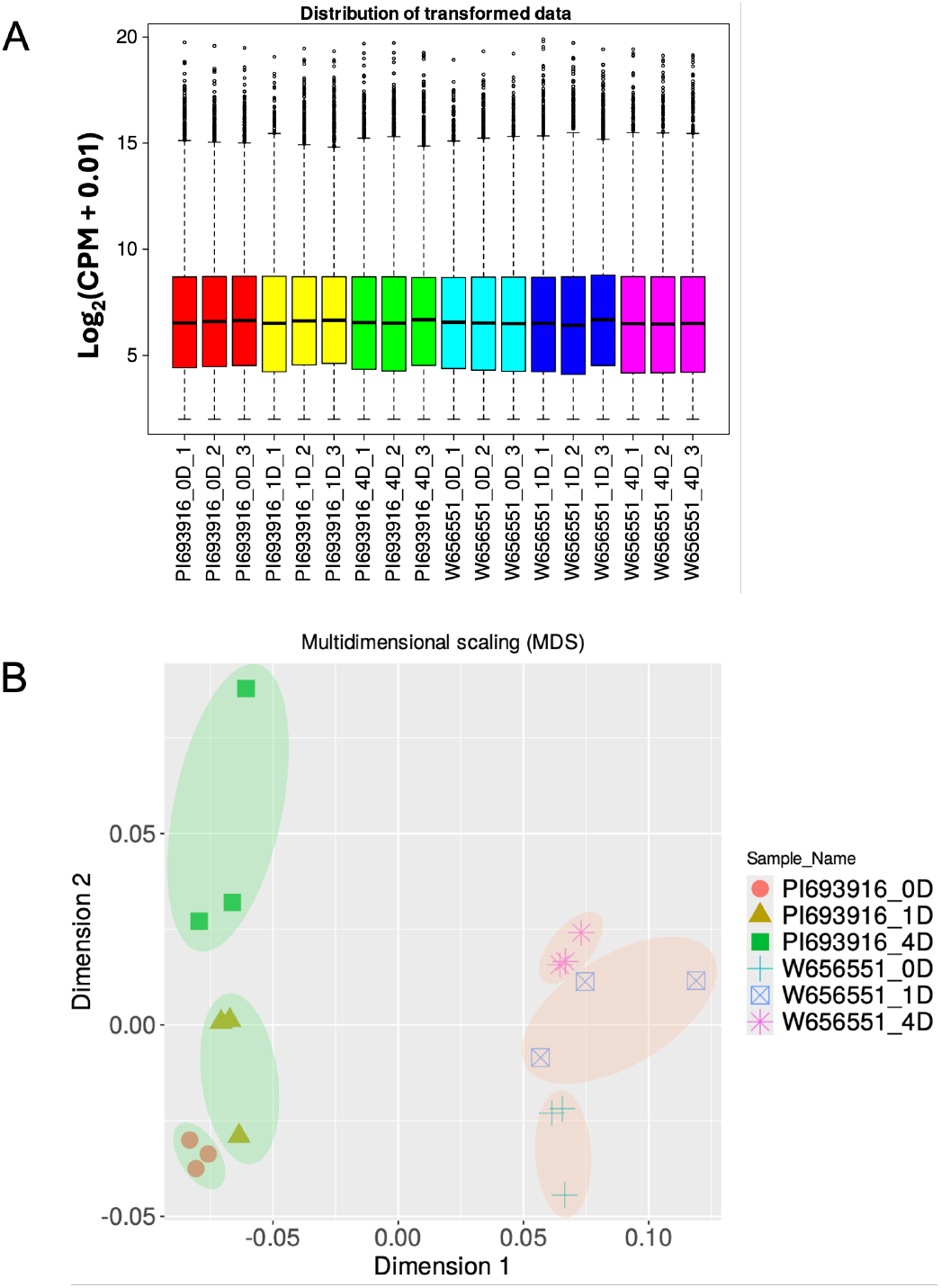
(A) Normalized gene expression in bluebunch wheatgrass leaf samples. Boxplots of counts per million (CPM) across all replicates for the tested samples. (B) Multidimensional scaling (MDS) plots illustrating expression variation across samples under different treatments. D0 denotes control samples with no treatment; D1 and D4 represent samples subjected to 160 mM NaCl for 1 day and 4 days, respectively.

**Figure S2.**
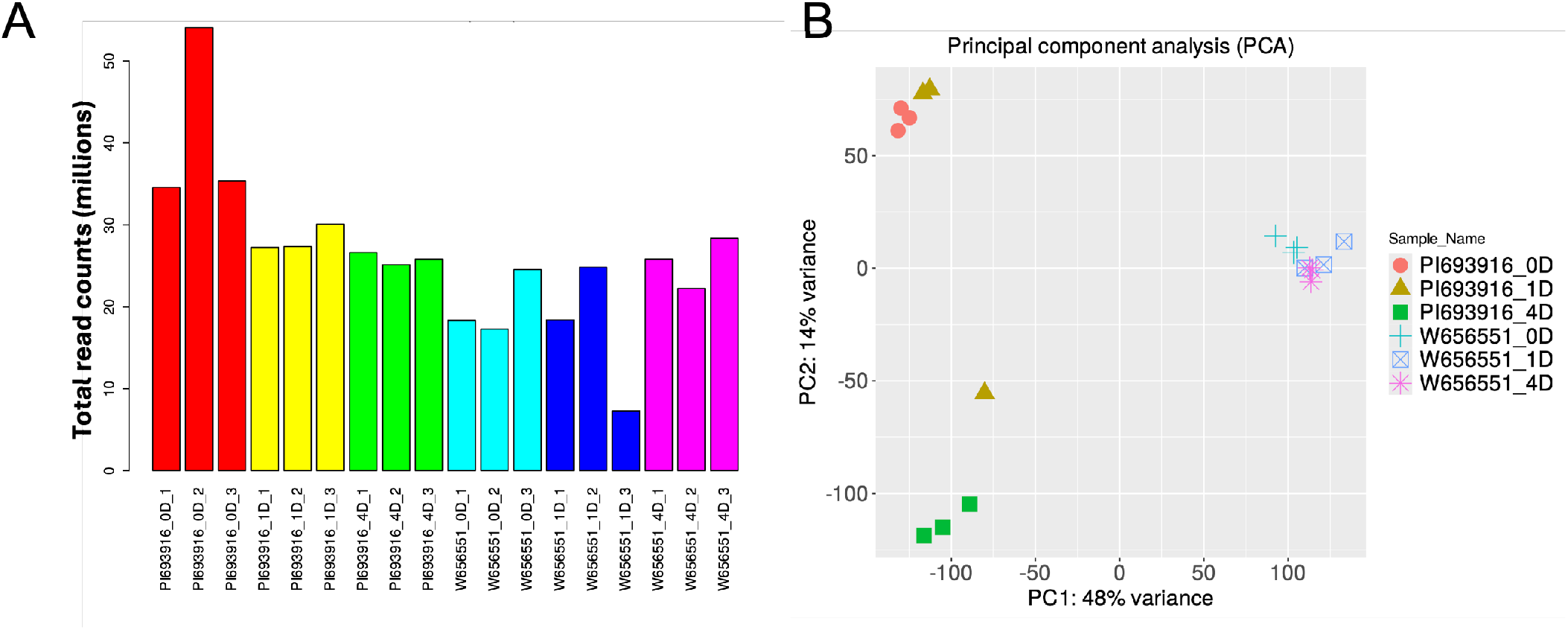
(A) Total read counts (in millions) for bluebunch wheatgrass leaf samples. (B) Principal component analysis of gene expression variation across samples under different treatments. 0D denotes control samples with no treatment; 1D and 4D represent samples subjected to 160 mM NaCl for 1 day and 4 days, respectively.

**Figure S3.**
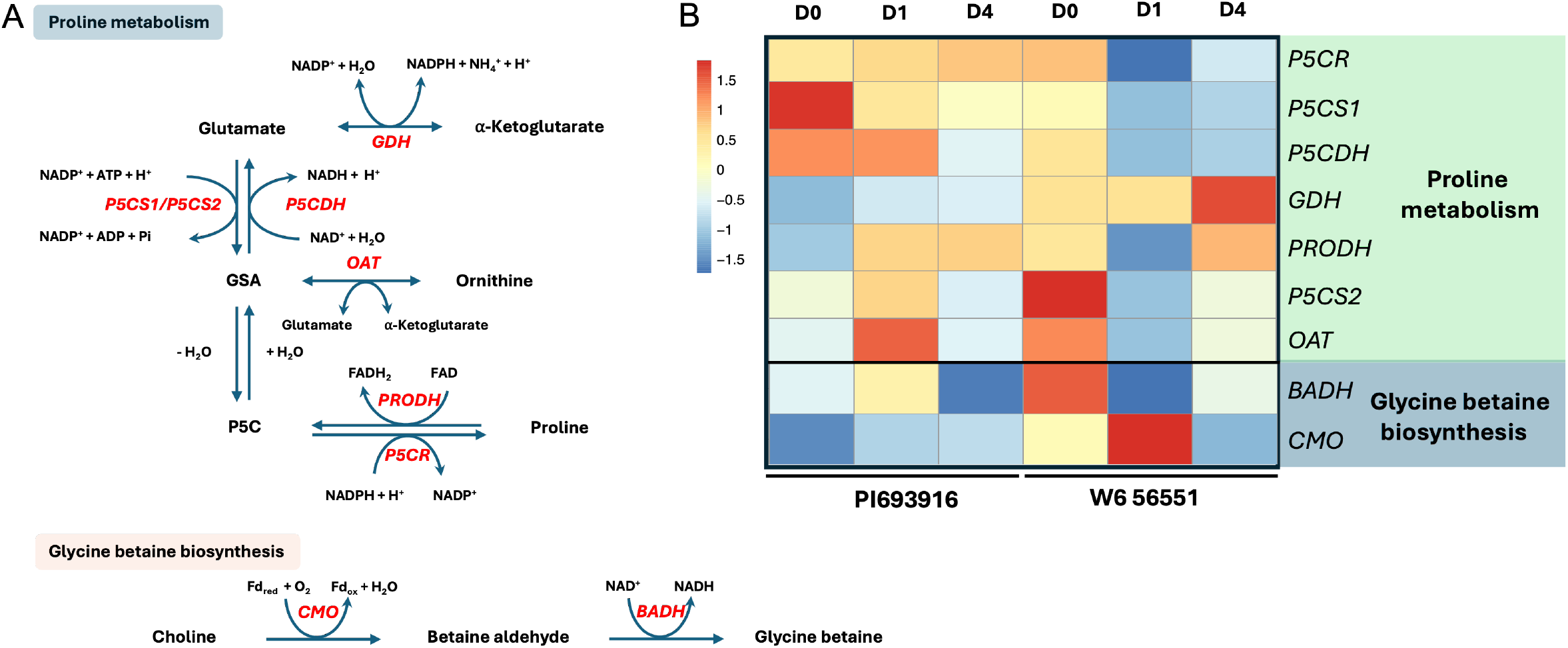
(A) Biosynthetic pathways of proline and glycine betaine in higher plants. *GDH*, glutamate dehydrogenase; *P5CS*, pyrroline-5-carboxylate synthetase; *P5CDH*, P5C dehydrogenase; *P5CR*, P5C reductase; *PRODH*, proline dehydrogenase; *OAT*, ornithine-*δ*-aminotransferase; *CMO*, choline monooxygenase; *BADH*, betaine aldehyde dehydrogenase. (B) Heatmap of expression of key genes involved in synthesis of Proline and Glycine betaine.

**Figure S4.**
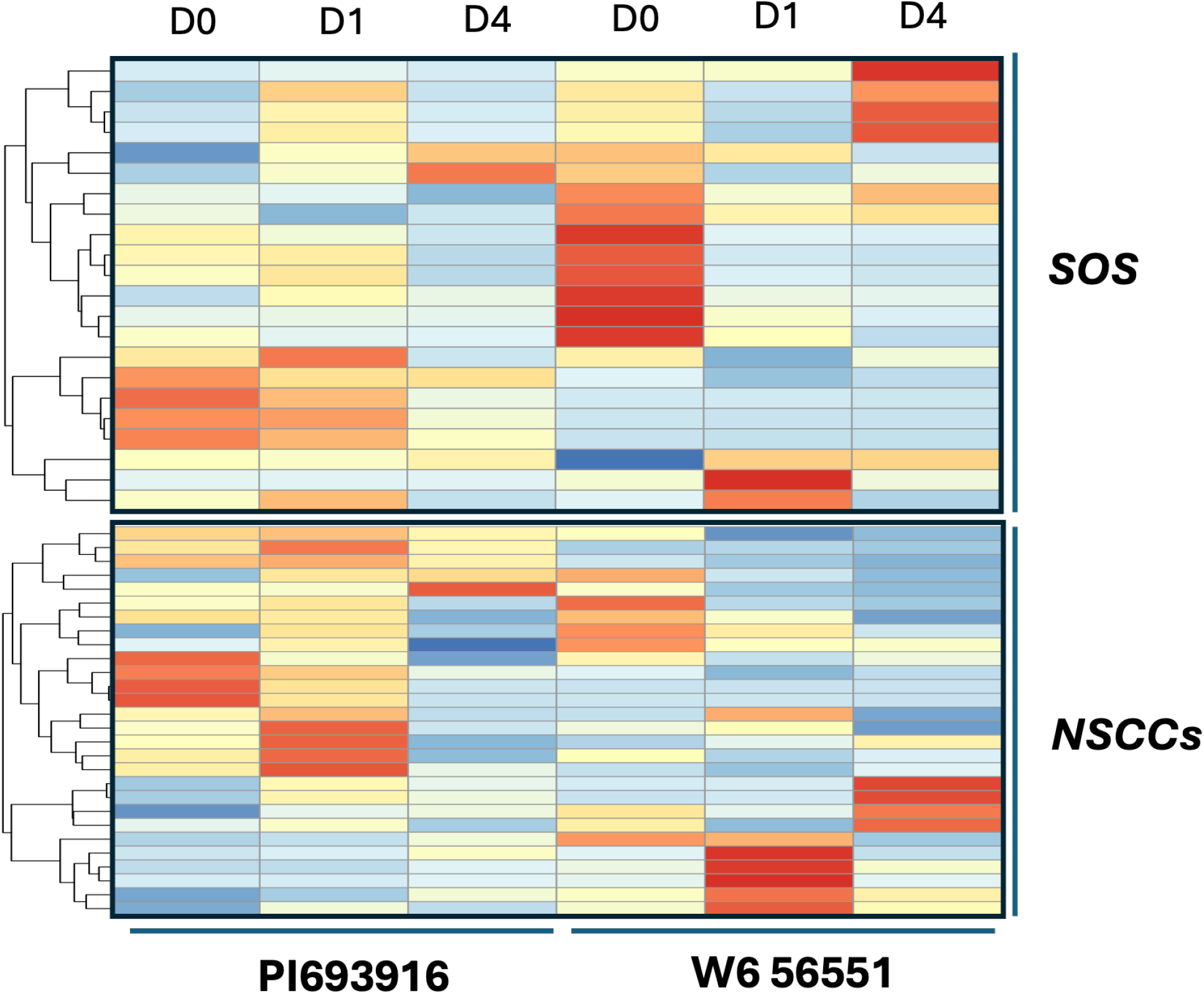
Heatmap of expression of *SOS, NSCCs* genes.

